# Optimization of Retinoid Detection in Cerebrospinal Fluid Using Liquid Chromatography Mass Spectrometry

**DOI:** 10.64898/2026.03.25.714054

**Authors:** Jeannette Brook, Xinran Tong, Alan Wong, Michal Weitman, Adrienne Boire, Naama Kanarek, Boryana Petrova

## Abstract

**Introduction:** Retinoids are bioactive vitamin A derivatives that regulate cellular differentiation and gene expression, yet their reliable quantification remains challenging due to low abundance, structural isomerism, and sensitivity to ionization conditions while handling.

**Objectives:** In this study, we performed a systematic optimization of liquid chromatography–mass spectrometry (LC-MS)-based detection of retinoids across tissues and biofluids.

**Methods:** Chromatographic separation, adduct formation, ionization parameters, fragmentation behavior, and extraction procedures were evaluated in an integrated workflow.

**Results:** Chromatographic conditions influenced not only retention time but also the ionic species detected, affecting precursor selection for MS² analysis. Retinoids exhibited compound-dependent responses to electrospray ionization and collision energy, requiring tailored acquisition parameters. Extraction experiments demonstrated differential recovery among retinoid classes and revealed matrix-dependent behavior, indicating that protocols used for tissues cannot be directly transferred to low-abundance biofluids. Using optimized conditions, retinoids were detected in mouse cerebrospinal fluid (CSF) at concentrations approaching the analytical detection limit, where MS² confirmation was necessary for reliable identification.

**Conclusion:** Together, our results provide a framework for reproducible retinoid profiling across biological matrices and enables comparative studies of retinoid biology in low-volume and low-abundance biofluids.

## 1 Introduction

Retinoids are bioactive metabolites derived from vitamin A that regulate gene expression and cellular differentiation through ligand-dependent nuclear receptor signaling. Retinoic acid (RA) functions as a transcriptional regulator across diverse tissues, where even small changes in local concentration can alter developmental and homeostatic programs(Das et al. 2014; Ghyselinck and Duester 2019). Dysregulation of retinoid metabolism and signaling has been implicated in multiple disease contexts, including cancer, where altered RA signaling affects differentiation and proliferation(Brown 2023), neurological disorders involving impaired neurogenesis and synaptic plasticity(Park et al. 2021), and metabolic diseases linked to disrupted lipid and energy homeostasis(Blaner 2019). These observations highlight that retinoid abundance is not merely descriptive but functionally determinant, motivating the need for accurate and reproducible quantification.

Endogenous retinoid pools are maintained through tightly controlled enzymatic interconversion between retinol, retinal, and retinoic acid, linking metabolic turnover directly to signaling activity(Neilson 2013). Because retinoid signaling depends on spatially and temporally restricted concentration gradients, analytical measurements must resolve small, localized differences rather than bulk systemic levels.

Reliable quantification is particularly challenging in two distinct but compounding contexts. First, biological sample volume is often limited, especially in applications such as cerebrospinal fluid (CSF), microdissected tissues, or small animal models. Second, retinoids are often present at low endogenous concentrations, frequently near the limits of detection, where local synthesis and degradation dominate over systemic abundance. These constraints necessitate analytical methods that are both highly sensitive and quantitatively robust.

Liquid chromatography–tandem mass spectrometry (LC–MS/MS) has enabled the detection of retinoic acid in small biological samples(Kane et al. 2005, 2008; Kane and Napoli 2010). However, retinoid levels in CSF and related compartments remain comparatively undercharacterized despite their relevance to neurological function and disease(Tabassi et al. 2005; Warner et al. 2007). Addressing these applications requires analytical workflows that maintain sensitivity while ensuring chemical specificity across structurally similar species.

Retinoid analysis is further complicated by challenges that arise at multiple stages of the analytical workflow. At the chemical level, the conjugated polyene structure permits geometric isomerization, while functional groups confer partial polarity and chemical reactivity. As a result, retinoids are susceptible to oxidation, light-induced degradation, and interconversion during sample handling – all are likely to happen while handling the samples(Furr 2004; Ioele et al. 2005). During sample preparation and extraction, these properties can lead to analyte loss, artifactual chemical conversion, and variable recovery depending on matrix composition.

At the level of chromatographic separation, structurally similar isomers require careful resolution to avoid co-elution, which directly impacts downstream identification. In the mass spectrometric stage, retinoids exhibit variable ionization efficiency, form multiple adduct species (e.g., sodium adducts) and undergo in-source fragmentation such as dehydration. These effects complicate precursor ion selection and fragmentation consistency, ultimately affecting quantitative reproducibility(Czuba et al. 2020). Additionally, matrix effects further modulate ionization behavior, particularly in complex samples such as tissues or biofluids.

Substantial methodological progress has been achieved using high-performance liquid chromatography (HPLC) and MS/MS for targeted retinoid analysis(Jones et al. 2015; Kane and Napoli 2010). However, most workflows optimize individual components, such as chromatographic separation or mass spectrometric detection, in isolation. In contrast, extraction efficiency, ionization behavior, and matrix effects are often treated separately and are not described in detail as part of the LC-MS protocol. As a result, differences in precursor ion selection, adduct formation, and extraction chemistry may therefore contribute to variability between studies without reflecting true biological differences(Czuba et al. 2020).

A systematic evaluation that integrates **sample preparation, chromatographic behavior, ion formation, and fragmentation** has not been comprehensively performed, particularly for low-volume biofluids. The extent to which chromatographic conditions influence adduct formation and precursor selection remains insufficiently characterized, and comparative assessment of extraction performance across tissues and biofluids is limited.

In the present study, we performed an integrated optimization of LC–MS-based retinoid analysis. We systematically evaluated chromatographic separation, characterized ionization and fragmentation behavior, compared extraction strategies across biological matrices, and assessed detection performance in CSF. This work provides a framework for reliable retinoid profiling in low-volume samples and aims to improve analytical consistency in studies of retinoid biology.

## 2 Methods

### 2.1 Retinoid standards preparation

Retinoid standards were weighed into small amber glass vials and solubilized in LC-MS grade methanol (A456-4, Optima™ Fisher Chemical) or ethanol (CAS No: Ethyl Alcohol 64-17-5) purged with N_2_ gas. Yellow light was used to prepare the standards and preserve stability. Standards were prepared at 1mM concentrations and stored at -80°C. We noted that the stability of the different retinoid standards in solution varied. For example, the 11-cis-retinol standard in solution was stable maximum up to a week. Freeze-thaw cycles of the retinoid standards were minimized, and fresh solutions were prepared at least every month. Standards are outlined in table M1:

**Table M1.**
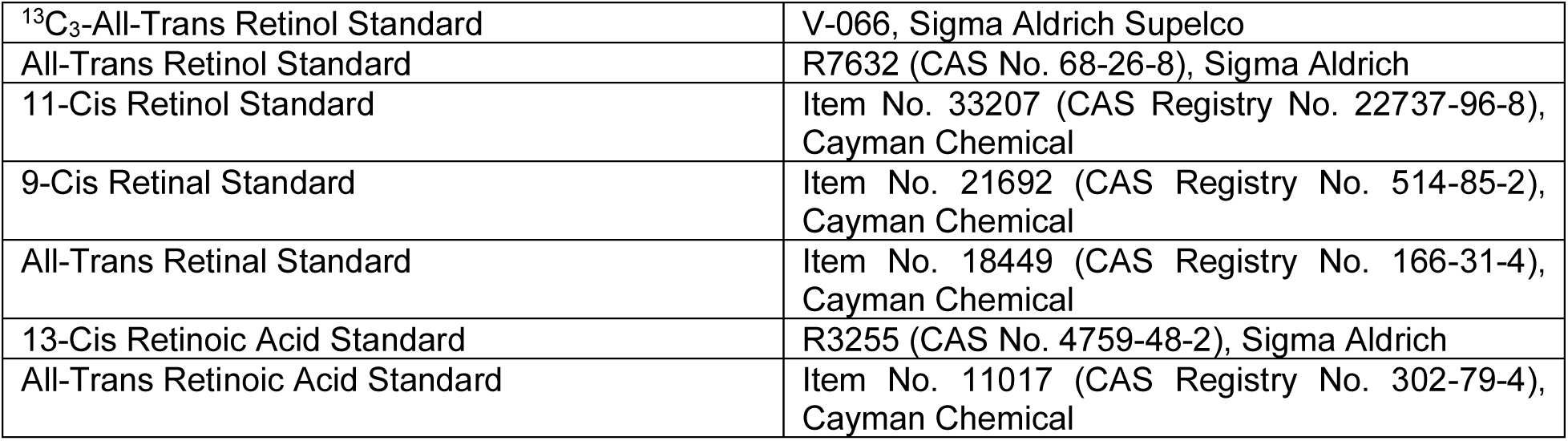
LC-MS retinoid standards.

|  |  |
| --- | --- |
| <sup>13</sup>C<sub>3</sub>-All-Trans Retinol Standard | V-066, Sigma Aldrich Supelco |
| All-Trans Retinol Standard | R7632 (CAS No. 68-26-8), Sigma Aldrich |
| 11-Cis Retinol Standard | Item No. 33207 (CAS Registry No. 22737-96-8), Cayman Chemical |
| 9-Cis Retinal Standard | Item No. 21692 (CAS Registry No. 514-85-2), Cayman Chemical |
| All-Trans Retinal Standard | Item No. 18449 (CAS Registry No. 166-31-4), Cayman Chemical |
| 13-Cis Retinoic Acid Standard | R3255 (CAS No. 4759-48-2), Sigma Aldrich |
| All-Trans Retinoic Acid Standard | Item No. 11017 (CAS Registry No. 302-79-4), Cayman Chemical |

### 2.2 Collection of tissues and biofluids for LCMS analysis or retinoids

Several different biological sample types were used to test optimal extraction and resuspension methods for retinoid metabolite detection.

#### 2.2.1 Cod liver and fish collection

Rice grain chunks of supermarket purchased cod liver or fresh and cooked salmon were separated into tubes and flash frozen for further analysis following procedures outlined below as for mouse CSF or liver.

#### 2.2.2 Mouse CSF and liver collection

The Institutional Animal Care and Use Committee (IACUC) at BCH approved all animal work carried out in this study (protocol #2068). NOD.CB17-*Prkdc*scid/NCrCrl (NOD-SCID) mice (originally purchased from Charles River Laboratories and maintained at the BCH mouse facility through in-house breeding) older than 3 months were anesthetized by intraperitoneal injection with ketamine and xylazine (120 mg/kg and 10mg/kg respectively). Cerebrospinal fluid (CSF) was collected from the cisterna magna(Courtney et al. 2025; Fame et al. 2024) using a glass micro-capillary needle (Fisher Scientific 21-176-2C) pulled using a dual-stage glass micropipette puller (Narishige PC-10). CSF collection was performed under yellow light to reduce oxidation of CSF metabolites. Following CSF collection, the anterior portion of the left lobe of liver was collected and snap-frozen in liquid nitrogen until further processing.

### 2.3 Extraction of tissues and biofluids for LCMS analysis of retinoid

Six different extraction methods were tested and specified as outlined in Table M2 Hexane and chloroform-based extractions were two-phase, while methanol-only based extraction was single phase. All extractions were performed on ice and under yellow light to minimize degradation and oxidation. Below further details are given per extraction type steps for unique extraction methods:

**Table M2.** Summary of examined methods for efficient extraction of retinoid metabolites from biological samples.

| Extraction components | Volumes used (μL) |
| --- | --- |
| 80% methanol | 300 μL |
| <u>2:3:1.5 chloroform:methanol:water</u> | 200 μL:300 μL:150 μL |
| <u>2:2:1.8 chloroform:methanol:water</u> | 200 μL:200 μL:180 μL |
| <u>2:1:0.5 chloroform:methanol:water</u> | 400 μL:200 μL:100 μL |
| hexanes, 0.025 M KOH in ethanol, 4 M HCl | 1 mL hexanes, 100 μL 0.025 M KOH in ethanol, 20-40 μL 4 M HCl |
| hexanes, acetonitrile, 4 M HCl | 1 mL hexanes, 100 μL ACN, 20-40 μL 4 M HCl |

#### 2.3.1 80% methanol extraction

To each sample 300 μL extraction buffer (80% methanol and 20% water, supplemented with ^13^C_3_-all-trans retinol (V-066, Sigma Aldrich Supelco) and 0.05% Butylated Hydroxytoluene (BHT) (PHR1117, Sigma Aldrich)) was added, samples were vortexed for 10 sec then centrifuged for 10 min at 18,000 g in a precooled benchtop centrifuge (Eppendorf 5424 R, Thermo 022431081). The supernatant was transferred to a new 1.5 mL Protein LoBind tube (Eppendorf Protein 022431081), dried on ice using nitrogen flow (Thermo Scientific, TS-18826) and resuspended in 50 μL 66% methanol (resuspension optimizations were performed separately, see results section).

#### 2.3.2 Methanol:chloroform:water based extractions

To each sample, appropriate volumes of chloroform (supplemented with UltimateSPLASH ONE, Avanti Polar Lipids, 330820), then methanol then water were added (respective volumes for each type of extraction are outlined in Table M2). Methanol was supplemented with ^13^C_3_-all-trans retinol and 0.05% BHT, while water was supplemented with isotopically labeled amino acids (MSK-A2-1.2, Cambridge Isotope Laboratories). Samples were vortexed for 10 sec then centrifuged for 10 min at 18,000 g in a precooled benchtop centrifuge (Eppendorf 5424 R, Thermo 022431081) to separate organic and aqueous phases. Each phase was transferred to a new 1.5 mL Protein LoBind tube (Eppendorf, 022431081) and dried using nitrogen flow (Thermo Scientific, TS-18826). Bottom (organic) phase was dried on ice, while top (aqueous) phase at room temperature (to speed up the process).

#### 2.3.3 Hexanes 0.025 M KOH in ethanol, 4 M HCl extraction

Extraction protocol was performed based on Kane & Napoli with the following modifications(Czuba et al. 2020; Kane and Napoli 2010). 100 μL of ethanol solution with 0.025M potassium hydroxide KOH, supplemented with ^13^C_3_-all-trans retinol and 0.05% BHT; 2) was added to each sample. Then 1 mL of hexanes (H303-1, Optima™ Fisher Chemical); 3) was added and samples were vortexed for 10 sec, centrifuged for 5 min at 18,000 g (Eppendorf 5424 R, Thermo 022431081) to separate organic and aqueous phases and the upper (organic) phase was transferred to a new 1.5 mL Protein LoBind tube (Eppendorf, 022431081) and stored on ice. 6μL of 4M hydrochloric acid (A144, Fisher Chemical) was added to the bottom aqueous layer, followed by 1mL hexanes, then vortexed for 10 sec and centrifuge for 5 min at 18,000 g to separate organic and aqueous phases a second round. The upper organic phase was transferred to a new 1.5 mL Protein LoBind tube (Eppendorf, 022431081). Finally, both organic phase extracts were dried on ice, using nitrogen flow (Thermo Scientific, TS-18826).

#### 2.3.4 Hexanes, ACN, 4M HCl extraction

Extraction protocol was performed based on Kane & Napoli with the following modifications(Czuba et al. 2020; Kane and Napoli 2010). 100μL acetonitrile supplemented with ^3^C_3_-all-trans retinol and 0.05% BHT was added to each sample followed by 6μL of 4M HCl and 1mL hexanes. Samples were vortexed for 10 sec then centrifuged for 5 min at 18,000 g (Eppendorf 5424 R, Thermo 022431081) to separate organic and aqueous phases. The upper organic layer was transferred to a new 1.5 mL Protein LoBind tube (Eppendorf, 022431081) and the upper organic phase was dried on ice, using nitrogen flow (Thermo Scientific, TS-18826).

Finally, as part of the sample preparation optimization, different reconstitution buffers were tested (supplemented with QReSS internal standards, Cambridge Isotope Laboratories, MSK-QreSS-KIT) as specified in Table M3 and further in the results section.

**Table M3.** Summary of examined methods for efficient resuspension of retinoid metabolites in biological material.

| # | resuspension components |
| --- | --- |
| #1 | 66% methanol |
| #2 | methanol |
| #3 | acetonitrile |
| #4 | 65% acetonitrile, 30% isopropanol, 5% water |

### 2.4 Liquid chromatography optimization conditions

Liquid chromatography optimization was performed in two steps: i. comparing different C18 columns or ii. Comparing different gradients on the column chosen from step 1. Standard dilutions were prepared in amber glass MS vials (VWR North American Cat. No. 10803-888A) to minimize oxidation during sample preparation. One μL of step wise dilutions (1mM to 100pM) was injected on one of three C18 chromatography columns tested (Table M4) and separated using the following condition C18_MetOH_2 outlined in Table M5; or on Ascentis® Express C18 column ran under the gradients outlined in Table M4. Columns were operated on a Vanquish™ Flex UHPLC System (Thermo Fisher Scientific, San Jose, CA, USA). The column oven and autosampler tray were held at 30 °C and 4 °C, respectively.

**Table M4.** LC-MS sample preparation materials and chromatography columns.

|  |  |
| --- | --- |
| Ascentis® Express C18 HPLC Column | 53825-U Sigma Aldrich Supelco, 2.7$\mu\text{m}$ particle size, L x I.D. 15 cm x 2.1 mm |
| Xselect Premier HSS T3 VanGuard FIT Column | 186009856, Waters™, 2.5 $\mu\text{m}$, 2.1 mm x 150 mm |
| Phenomenex C18 LC Column | 00F-4462-Y0, Phenomenex Kinetex® 2.6$\mu\text{m}$ C18 500 Å, 100 x 4.6 mm |

**Table M5.** Chromatography gradients tested for LC optimization using retinoid standards on Ascentis® Express C18 HPLC column.

| Method | Weaker Eluent (A) | Stronger Eluent (B) | Gradient conditions | Flow Rate |
| --- | --- | --- | --- | --- |
| C18-FA-1 | LC-MS Water<br>0.1% Formic Acid | ACN<br>0.1% Formic Acid | 0-2 min: 5% B<br>2-10 min: 5%-36% B<br>10-10.1 min: 36%-95% B<br>10.1-14 min: hold 95% B<br>14 -14.1 min: 5% B<br>14.1 -18 min: 5% B | 0.250 $\mu\text{L}/\text{min}$ |
| C18-FA-2 | LC-MS Water<br>0.1% Formic Acid | ACN<br>0.1% Formic Acid | 0-2 min: 5% B<br>2-8 min: 5%-80% B<br>8-12 min: 80%-95% B<br>12-17 min: hold 95% B<br>17-17.1 min: 95%-5% B<br>17.1-22 min: 5% B | 0.250 $\mu\text{L}/\text{min}$ |
| C18-FA-3 | LC-MS Water<br>0.1% Formic Acid | LC-MS ACN<br>0.1% Formic Acid | 0-1 min: hold 10% B<br>1-11 min: 10%-65% B<br>11-14 min: hold 65% B<br>14-14.2 min: 65%-95%<br>14.2-17 min: hold 95% B<br>17-17.1 min: 95%-10% B<br>17.1-22 min: hold 10% B | 0.250<br>μL/min |
| C18-Lipid<br>(Cantor et al.,<br>2017) | LC-MS Water<br>0.1% Formic Acid,<br>10 mM ammonium<br>formate | LC-MS ACN<br>0.1% Formic Acid,<br>10 mM ammonium<br>formate | 0-1.5 min: hold 32% B<br>1.5-4 min: 32%-45% B<br>4-5 min: 45%-52% B<br>5-8 min: 52%-58% B<br>8-11 min: 58%-66% B<br>11-14 min: 66%-70% B<br>14-18 min: 70%-75% B<br>18-21 min: 75%-97% B<br>21-25 min: hold 97% B<br>25-25.1 min: 92%-32% B<br>25.1-30 min: hold 32% B | 0.250<br>μL/min |
| C18-MetOH-1<br>(Doyle, et al.,<br>2017) | LC-MS Water<br>0.1% Formic Acid | LC-MS Methanol | 0-2 min: hold 50% B<br>2-10 min: 50%-98% B<br>10-14 min: hold 98% B<br>14-14.1 min: 98%-50% B<br>14.1-18 min: hold 50% B | 0.250<br>μL/min |
| C18-MetOH-2 | LC-MS Water<br>0.1% Formic Acid | LC-MS Methanol | 0-2 min: hold 75% B<br>2-10 min: 75%-98% B<br>10-14 min: hold 98% B<br>14-14.1 min: 98%-75% B<br>14.1-18 min: hold 75% B | 0.250<br>μL/min |

#### 2.4.1 Optimal liquid chromatography conditions

Following sample reconstitution, 5-8 μL were injected into an Ascentis® Express C18 HPLC column (15 cm x 2.1 mm, particle size 2.7 μm; Sigma Aldrich) operated on a Vanquish™ Flex UHPLC System (Thermo Fisher Scientific, San Jose, CA, USA). The column oven and autosampler tray were held at 30 °C and 4 °C, respectively. The following conditions were used to achieve chromatographic separation: buffer A: water with 0.1% formic acid; buffer B: methanol. The chromatographic gradient was run at a flow rate of 0.250 ml min^−1^ as follows: 0–2 min: gradient was held at 75% B; 2–14 min: linear gradient of 75% to 98% B; 14.1–18.0 min: gradient was returned to 75% B.

### 2.5 Mass spectrometer operating conditions for retinoid detection

MS data acquisition was performed using a QExactive benchtop orbitrap mass spectrometer equipped with an Ion Max source and a HESI II probe (Thermo Fisher Scientific, San Jose, CA, USA). The mass spectrometer was operated in full-scan, positive ionization mode using one narrow-range scan: 260–310 m/z from 8-12 min with resolution at 70,000, and AGC target at 1 × 10^6^, and the maximum injection time at 100 ms. HESI settings tested are further specified in Table M6. Optimal HESI methods were: Sheath gas flow rate: 40 psi; Aux gas flow rate: 10 a.u.; Sweep gas: 0 a.u.; Spray voltage: 3.5 kV (pos); Capillary temperature 300 °C; S-lens RF level: 50 a.u.; Aux gas heater temp: 350 °C.

**Table M6.** HESI settings tested for MS optimization using retinoid standards.

| # | S-lens | capillary temperature (C) | aux gas temperature (C) | sheath gas flow rate | aux gas flow rate | sweep gas flow rate | spray voltage pos (kV) |
| --- | --- | --- | --- | --- | --- | --- | --- |
| 1 | 40 | 300 | 350 | 40 | 10 | 0 | 3.5 |
| 2 | 45 | 300 | 350 | 40 | 10 | 0 | 3.5 |
| 3 | 50 | 300 | 350 | 40 | 10 | 0 | 3.5 |
| 4 | 55 | 300 | 350 | 40 | 10 | 0 | 3.5 |
| 5 | 60 | 300 | 350 | 40 | 10 | 0 | 3.5 |
| 6 | 65 | 300 | 350 | 40 | 10 | 0 | 3.5 |
| 7 | 70 | 300 | 350 | 40 | 10 | 0 | 3.5 |
| 8 | 50 | 250 | 300 | 40 | 10 | 0 | 3.5 |
| 9 | 50 | 300 | 350 | 40 | 10 | 0 | 3.5 |
| 10 | 50 | 320 | 350 | 40 | 10 | 0 | 3.5 |
| 11 | 50 | 350 | 400 | 40 | 10 | 0 | 3.5 |
| 12 | 50 | 400 | 450 | 40 | 10 | 0 | 3.5 |
| 13 | 50 | 450 | 450 | 40 | 10 | 0 | 3.5 |

### 2.6 Metabolomics data analysis for retinoid detection

For retinoid standards relative quantification was performed with TraceFinder 5.1 (Thermo Fisher Scientific, Waltham, MA, USA) using a 5 ppm mass tolerance. After normalization to the internal standard, areas under the curve were plotted in GraphPad Prism 10. For biological samples relative quantification of extracted metabolites was performed with TraceFinder 5.1 (Thermo Fisher Scientific, Waltham, MA, USA) using a 5 ppm mass tolerance and referencing in-house chemical standards. Pooled samples and fractional dilutions were prepared as quality controls and injected at the beginning and end of each run. In addition, pooled samples were interspersed throughout the run to control for technical drift in signal quality as well as to serve to assess the coefficient of variability (CV) for each metabolite.

Data normalization was performed in two steps. (1) Peak areas of the internal standard ¹³C₃-all-trans retinol, included in the extraction buffer, were integrated for each sample, mean-centered, and averaged across samples. Individual samples were divided by this factor to correct for technical variability arising from MS signal fluctuation, pipetting, and injection errors (typically within ∼10%). (2) Normalization for biological material was based on endogenous metabolites. For each metabolite, coefficients of variation (CV; based on pooled sample reinjections) and coefficients of determination (R²; based on linear dilutions of pooled samples) were calculated. Metabolites with CV < 30% and R² > 0.95 were mean-centered and averaged to generate a biological normalization factor, and metabolite peak areas were divided by this factor to account for global differences in metabolite abundance between samples.

For comparisons across biofluids, normalization was additionally performed relative to the extracted sample volume. When different sample volumes were processed, extraction and reconstitution solvent volumes were proportionally adjusted to maintain a consistent matrix concentration. For tissue samples, sample weight was used as an additional normalization factor.

**Table M7.** LC-MS solvents, reagents, and standards.

| Item | Catalogue Information |
| --- | --- |
| LC-MS Water | W6-4, Optima™ Fisher Chemical |
| LC-MS Methanol | A456-4, Optima™ Fisher Chemical |
| LC-MS Acetonitrile | A955-4, Optima™ Fisher Chemical |
| LC-MS Isopropanol | A461-2, Optima™ Fisher Chemical |
| LC-MS Chloroform | C606SK-1, Optima™ Fisher Chemical |
| LC-MS Hexanes | H303-1, Optima™ Fisher Chemical |
| LC-MS Formic Acid | 00940, LiChropur™ Sigma Aldrich |
| LC-MS Ammonium Carbonate | 37999, Sigma Aldrich |
| LC-MS Ammonium Hydroxide | 5.43830, Sigma Aldrich |
| Potassium Hydroxide (KOH) | P5958, Honeywell (1310-58-3) |
| Ethyl Alcohol 200 Proof | 111000200, Pharmco by Greenfield Global (CAS No: Ethyl Alcohol 64-17-5) |
| Hydrochloric Acid (HCl) | A144, Fisher Chemical (CAS 7647-01-0) |
| UltimateSPLASH™ ONE Internal Standard for Lipidomic Analysis | 330820, Avanti® Polar Lipids |
| Metabolomics QReSS™ Standard Mix | MSK-QReSS-KIT, Cambridge Isotope Laboratories |
| Metabolomics Amino Acid Mix | MSK-A2-1.2, Cambridge Isotope Laboratories |
| Butylated Hydroxytoluene | PHR1117 (CAS-No: 128-37-0), Sigma Aldrich |

**Table M9.**
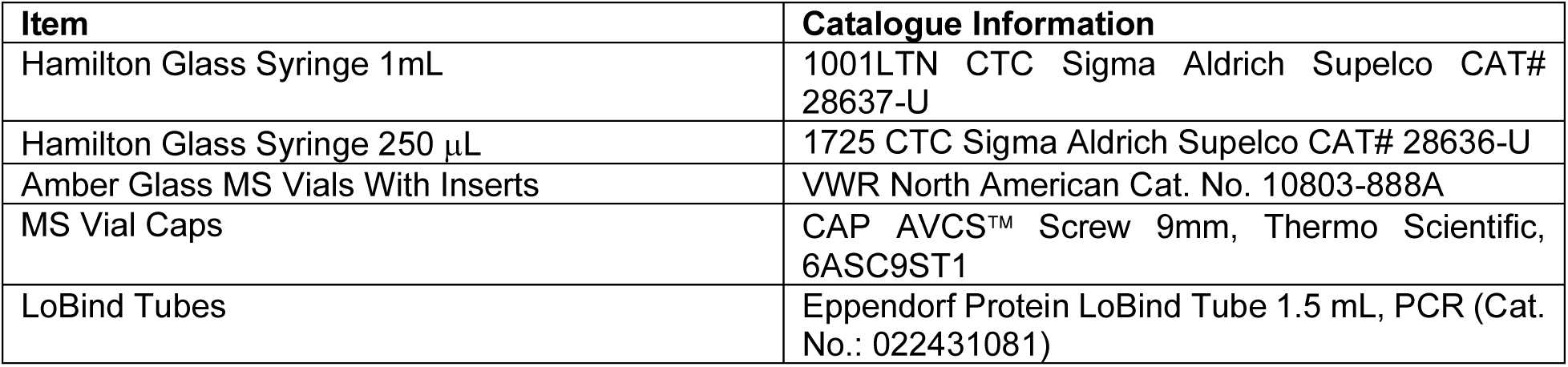
LC-MS sample preparation materials.

| Item | Catalogue Information |
| --- | --- |
| Hamilton Glass Syringe 1mL | 1001LTN CTC Sigma Aldrich Supelco CAT# 28637-U |
| Hamilton Glass Syringe 250 µL | 1725 CTC Sigma Aldrich Supelco CAT# 28636-U |
| Amber Glass MS Vials With Inserts | VWR North American Cat. No. 10803-888A |
| MS Vial Caps | CAP AVCS™ Screw 9mm, Thermo Scientific, 6ASC9ST1 |
| LoBind Tubes | Eppendorf Protein LoBind Tube 1.5 mL, PCR (Cat. No.: 022431081) |

## 3 Results

### 3.1 Optimization of liquid chromatography conditions for retinoid detection

To establish robust chromatographic conditions for retinoid detection, we initially focused on all-trans retinol and 13-cis retinoic acid as representative analytes covering distinct chemical classes within retinoid metabolism (Fig. 1A). Method development was guided by systematic evaluation of reversed-phase chromatographic parameters, including stationary phase, mobile phase composition, and gradient design (schematic overview in Fig. S1A).

**Figure 1.**
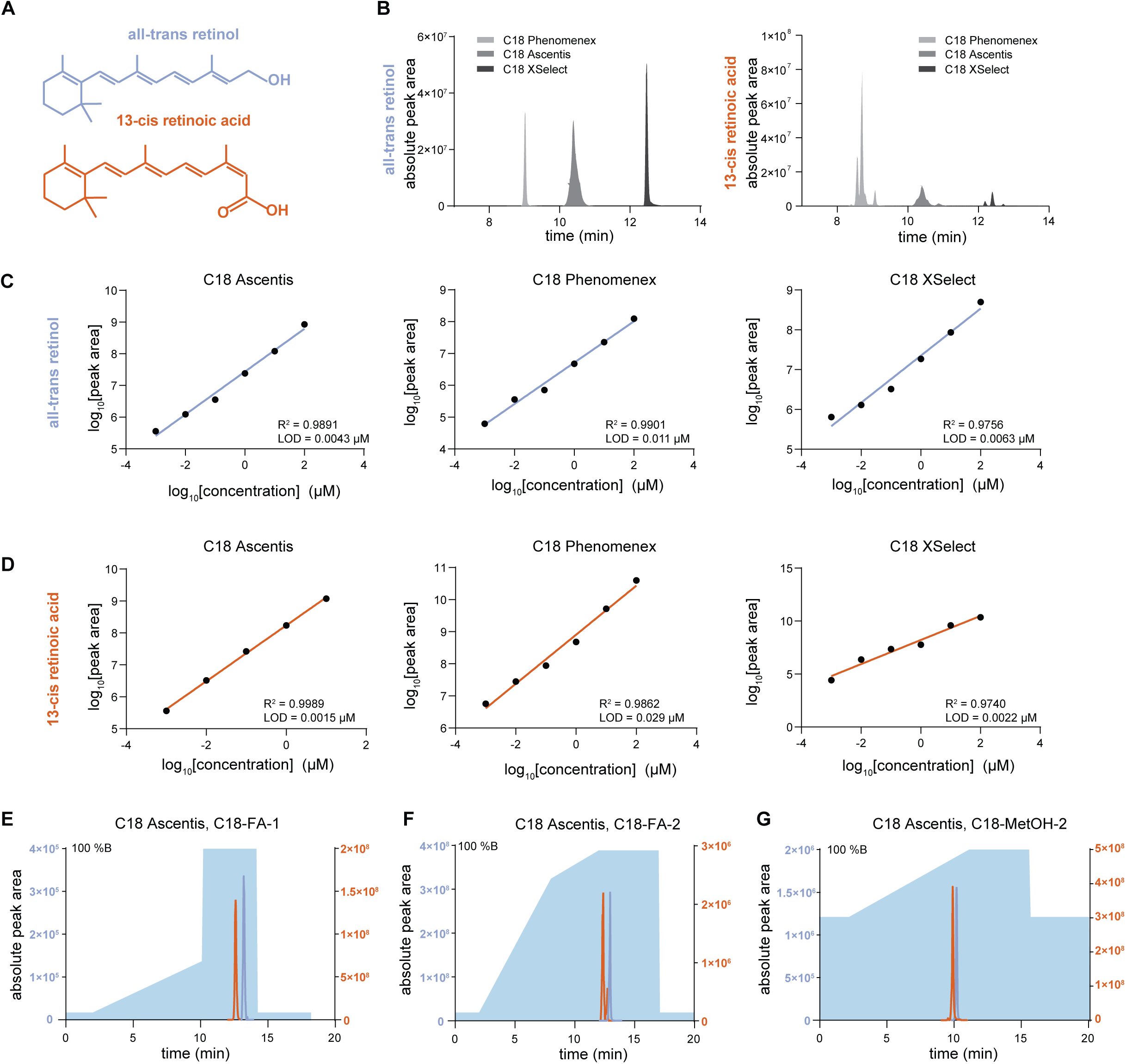
Optimization of liquid-chromatography method for all-trans retinol and 13-cis retinoic acid detection. **(A)** Chemical structures of all-trans retinol and 13-cis retinoic acid. **(B))** Extracted ion chromatograms for all-trans retinol (m/z 269.2264) (left) and 13-cis retinoic acid (m/z 301.2162) (right) using the chromatographic gradient in **(G)** on three different C18 columns: C18 Ascentis, C18 Phenomenex (2.6 μm C18 500 Å particle size, 100 x 4.6 mm), and C18 XSelect (2.5 μm particle size, 2.1 mm x 150 mm). Absolute intensity is plotted to allow for inter-column comparisons. **(C, D)** Limit of detection (LOD) and linearity for all-trans retinol and 13-cis retinoic acid standards (diluted in methanol) on three different C18 columns (as in **E**). Shown is representative experiment. A representative experiment (from minimum of 4 independent experiments) is shown. Linear regression analysis was performed, and R^2^-values are indicated. **(E, F, G)** Extracted ion chromatograms for all-trans retinol (m/z 269.2264) and 13-cis retinoic acid (m/z 301.2162) standards for three different chromatography gradients run on Ascentis Express C18 HPLC Column (2.7 μm particle size, 15 cm x 2.1 mm). Mobile phase A for all three gradients was 0.1% formic acid in water; mobile phase B was 0.1% formic acid in acetonitrile **(E, F)** or methanol **(G)**. Further details on each gradient can be found in methods section.

Three C18 columns with differing particle sizes and surface chemistries were compared under identical chromatographic gradients to assess signal intensity, peak shape, and retention behavior (Fig. 1B). These were: Ascentis Express, particle size 2.7 µm; Xselect HSS T3, particle size 2.5 µm; Phenomenex C18, particle size 2.6 µm (for further details see Methods). While both analytes were detectable on all tested columns, pronounced differences in detection between columns were observed. Each C18 column displayed distinct advantages in terms of peak shape and peak intensity for both compounds. Analysis on the Phenomenex and XSelect columns produced slightly narrower peaks than the Ascentis column but revealed multiple partially resolved peaks for 13-cis retinoic acid, suggesting enhanced isomeric separation or increased secondary structure interactions under these conditions.

Next, analytical sensitivity and linearity were systematically evaluated for all-trans retinol and 13-cis retinoic acid across the tested chromatographic platforms. Limit of detection (LOD) and linear dynamic ranges were determined using serial dilutions of analytical standards, with reproducibility assessed across independent experiments (Fig. 1C–D). Among the evaluated columns, the Ascentis and XSelect columns outperformed Phenomenex, achieving limits of detection of 0.0015 µM and 0.0022 µM for 13-cis retinoic acid and 0.0043 µM and 0.0063 µM for all-trans retinol respectively. Linear responses also varied across the tested concentration ranges for both analytes, with high coefficients of determination, indicating suitability for quantitative applications. These results supported the selection of the Ascentis column as the preferred stationary phase for retinoid analysis, achieving low LOD, broad linearity, and robust sensitivity for both analytes.

To further improve chromatographic performance, multiple gradient profiles and organic mobile phase compositions were evaluated using the Ascentis C18 column. Gradients employing either acetonitrile or methanol as the organic modifier were compared with respect to retention time, peak width, signal intensity, and separation between analytes (Fig. 1E–G). Methanol-based gradients consistently resulted in improved peak shape and more stable retention behavior for both all-trans retinol and 13-cis retinoic acid. Among the tested conditions, the C18-MetOH-2 gradient achieved elution of both analytes in the mid-gradient range while maintaining narrow peak widths and robust signal intensity. This gradient was therefore selected for all subsequent analyses.

Using the optimized chromatographic conditions, method performance was next evaluated across a broader panel of retinoid standards encompassing structural isomers or retinoid acids, aldehydes, and alcohols (Fig. S1B). LODs and linearity were assessed individually for each compound (Fig. S1C, S1E). All tested retinoids were detectable within the low micromolar range, although sensitivity varied substantially between compounds. All-trans retinal represented the least sensitive analyte under these conditions, with a limit of detection of 0.027 µM, highlighting compound-specific differences in ionization efficiency and chromatographic behavior.

Chromatographic separation of structurally related retinoid isomers was evaluated under the optimized LC conditions. Effective separation was achieved between all-trans retinoic acid and 13-cis retinoic acid, as reflected by distinct retention times and well-resolved peak shapes (Fig. S1D). In contrast, 9-cis retinal and all-trans retinal were not chromatographically separable under these conditions. Additionally, 11-cis retinol consistently exhibited a double peak, suggesting the presence of additional isomeric species or conformers, an observation further supported by MS^1^ adduct analysis, as further discussed below (Fig. S2F). These findings delineate the resolving power and limitations of the optimized LC method with respect to retinoid isomer separation.

### 3.2 Characterization of retention time and adduct formation for retinoid standards

Following chromatographic optimization, MS^1^ adduct formation was characterized for all-trans retinol and 13-cis retinoic acid under the selected LC conditions using the Ascentis column. Full-scan MS^1^ spectra revealed distinct and reproducible adduct profiles for both compounds (Fig. 2A). All-trans retinol was detected predominantly as the dehydrated protonated ion [M+H-H_2_O]^+^, whereas 13-cis retinoic acid was detected as a mixture of protonated and sodium-containing species, including [M+H]^+^, [M+Na]^+^, and a secondary sodium-containing adduct [M-H +Na_2_]^+^. ^13^C_3_ all-trans retinol behaved like the unlabeled form (Fig S2A). Based on signal intensity and consistency, [M+H-H_2_O]^+^ was selected for all-trans retinol and [M+H]^+^ for 13-cis retinoic acid in subsequent analyses.

**Figure 2.**
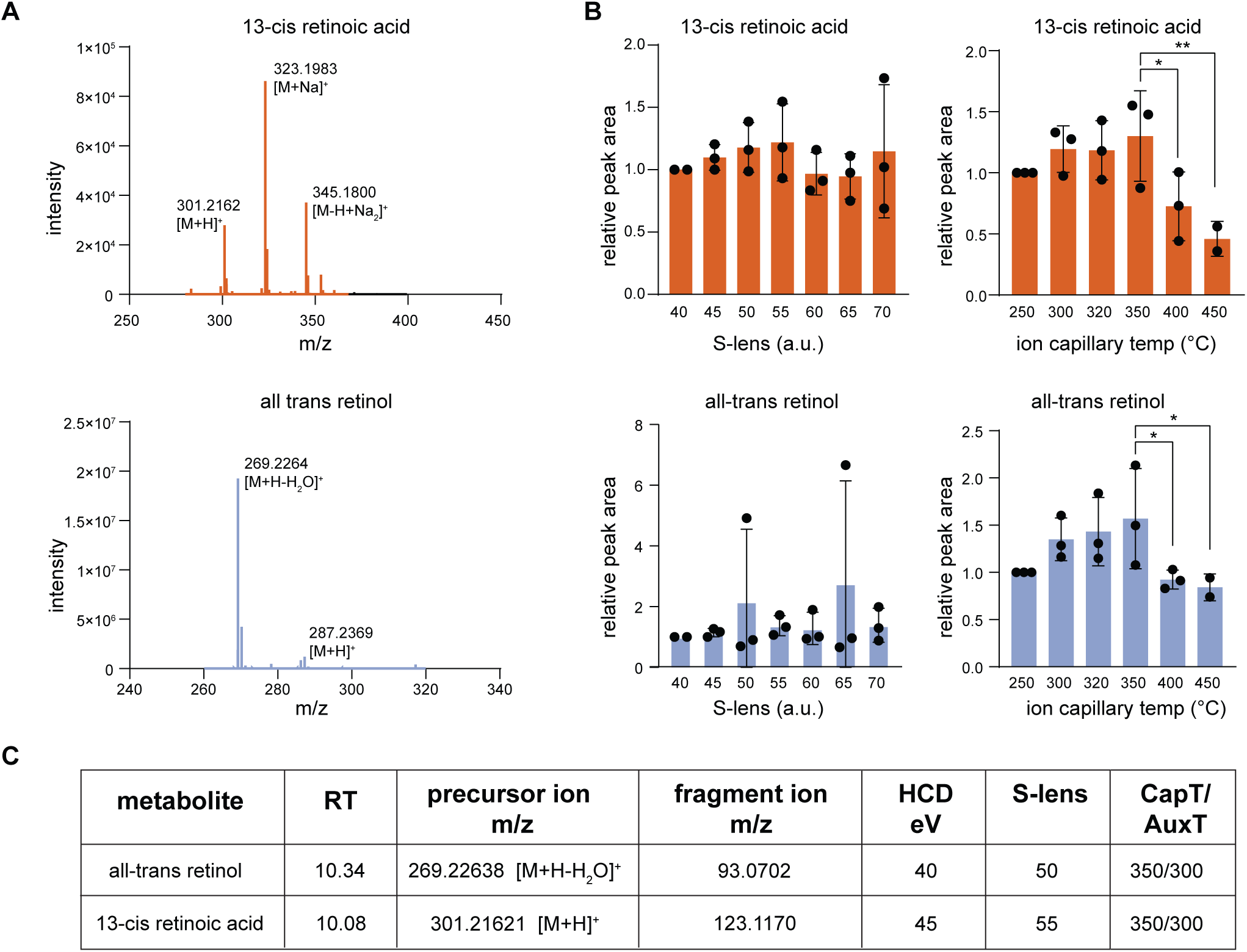
Optimization of mass spectrometry method and ionization parameters for the detection of all-trans retinol and 13-cis retinoic acid. (A) Predominant annotated adducts for all-trans retinol: [M+H-H_2_O]^+^ and [M+H]^+^ and 13-cis retinoic acid: [M+H]^+^, [M+Na]^+^, and [M-H+Na_2_]^+^ are indicated as observed from MS^1^ spectra. (B) HESI parameter optimization of S-lens (a.u.) and ion capillary temperature (°C) on an Orbitrap mass spectrometer. Bar plots show mean ± standard deviation from three experiments, representing chromatographic peak areas of retinoid metabolites under varying S-lens voltages and ion transfer capillary temperatures. p-values were obtained by one-way ANOVA with Šídák’s correction for multiple comparisons: * p<0.05, ** p<0.01. (C) Summary of retention times, precursor m/z values, and unique fragment ions used for MS² quantification of retinoid metabolites on the C18 Ascentis column with gradient 2.

To assess the influence of chromatographic conditions on adduct formation, MS^1^ spectra of 13-cis retinoic acid were compared across the three evaluated C18 columns (Fig. S2B–D). In all cases, multiple partially resolved peaks corresponding to the same analytical standard were observed, with column-dependent differences in both retention behavior and relative adduct abundance. On the Ascentis column, the predominant peak (RT 10.40 min) exhibited a balanced distribution of [M+H]^+^, [M+Na]^+^, and [M-H +Na_2_]^+^, whereas a later eluting peak (RT 10.86 min) was dominated by the protonated species. A similar pattern was observed on the XSelect column, where early and main peaks displayed mixed adduct distributions, while later eluting peaks showed increased dominance of [M+H]^+^. In contrast, analysis on the Phenomenex column yielded a more uniform adduct profile across all observed peaks, with [M+H]^+^ as the dominant species. Together, these observations demonstrate that chromatographic conditions substantially influence both retention behavior and MS1 adduct formation for retinoids. This variability underscores the importance of column selection and adduct characterization for robust precursor ion selection and quantitative analysis.

MS^1^ adduct characterization was next extended to additional retinoid standards analyzed under the optimized Ascentis chromatographic conditions, including all-trans retinoic acid, 9-cis retinal, and 11-cis retinol (Fig. S2E–G). All-trans retinoic acid was detected primarily as the protonated molecular ion [M+H]+, with a secondary methanol adduct present at lower abundance. In contrast, 9-cis retinal exhibited a broader adduct profile, with [M+H]+ and [M+Na]+ as dominant species, while two additional lower-abundance adducts could not be confidently annotated. In the case of 11-cis retinol, two chromatographically distinct peaks were consistently observed; however, both peaks showed highly similar adduct distributions, suggesting comparable ionization behavior despite chromatographic separation.

Together, these results establish a chromatographically and ionization-consistent framework for precursor ion selection, providing the basis for systematic optimization of MS source and fragmentation parameters described in the next section.

### 3.3 Optimization of mass spectrometry and ionization parameters

To maximize detection sensitivity for retinoids under the optimized chromatographic conditions, key heated electrospray ionization (HESI) source parameters were systematically evaluated (Fig. S3A), with a focus on S-lens voltage and ion transfer capillary temperature. Variation of the S-lens voltage resulted in reproducible, compound-dependent effects on chromatographic peak areas (Fig. 2B; Fig. S3B). Peak areas for all-trans retinol and 13-cis retinoic acid remained largely unchanged across the tested S-lens range, whereas increasing S-lens voltage generally enhanced sensitivity for all-trans retinoic acid, all-trans retinal, 9-cis retinal, and 11-cis retinol. In contrast, ion transfer capillary temperature had a pronounced and compound-specific impact on retinoid signal intensity (Fig. 2B; Fig. S3C). For all-trans retinol and 13-cis retinoic acid, intermediate temperatures yielded the highest and most stable peak areas, while temperatures above 350 °C strongly suppressed signal intensity. Increasing temperature reduced signal intensity for all-trans retinal and 9-cis retinal, whereas signals for all-trans retinoic acid and 11-cis retinol were comparatively unaffected across the tested range. Based on these combined observations, a single set of HESI parameters was selected, S-lens voltage 50 a.u. and ion transfer capillary temperature 300 °C, as a middle value that provided robust signal intensity across all tested retinoids and was fixed for all subsequent analyses.

To enable MS²-based quantification, fragmentation of all-trans retinol and 13-cis retinoic acid was evaluated at HCD energies of 20, 40, and 80 eV (Fig. S3D–E). At low collision energy (20 eV), fragmentation was limited and precursor ions dominated the spectra. At high collision energy (80 eV), fragmentation efficiency increased but resulted in reduced intensity of individual diagnostic ions and increased spectral complexity. Intermediate collision energies yielded the most robust and selective fragment ions for both analytes. For all-trans retinol, HCD 40 eV produced a dominant fragment at m/z 93.0702, whereas for 13-cis retinoic acid, HCD 45 eV maximized the abundance of the diagnostic fragment at m/z 123.1170. These conditions were therefore selected for PRM-based quantification, and the corresponding retention times, precursor ions, fragment ions, and collision energies are summarized in Fig. 2C.

To validate retinoid identification in biological samples, MS² spectra acquired from endogenous signals were directly compared to spectra obtained from in-house analytical standards using mirror plot analysis (Fig. S4). Comparisons were performed for all retinoids detected in biological matrices, using the optimized chromatographic and MS² acquisition parameters. For each compound, MS² spectra from biological samples closely matched those of the corresponding standards, with concordant retention times and highly similar fragmentation patterns (>97% overlap). Diagnostic fragment ions observed in standards were consistently present in biological spectra, and their relative intensities were preserved across matrices.

### 3.4 Optimization of sample preparation for tissue-based retinoid analysis

To optimize sample preparation for tissue-based retinoid analysis, we first compared biphasic extraction strategies using mouse liver and eye tissue, focusing on recovery and phase distribution of all-trans retinol and 13-cis retinoic acid (Fig. 3A). Given the well-known susceptibility of retinoids to light-induced degradation and oxidation, all extraction procedures were performed on ice and under yellow light to preserve analyte integrity—an essential condition for reproducible quantification. Two extraction approaches were evaluated: a chloroform/methanol/water-based extraction (2:2:1.8) and a two-step hexanes-based extraction. Distinct compound-dependent partitioning behaviors were observed. All-trans retinol was predominantly recovered in the first hexanes phase and in the bottom chloroform phase whereas 13-cis retinoic acid showed equal partition between both hexanes phases and a slightly higher abundance in the top phase of the chloroform/methanol/water extraction.

**Figure 3.**
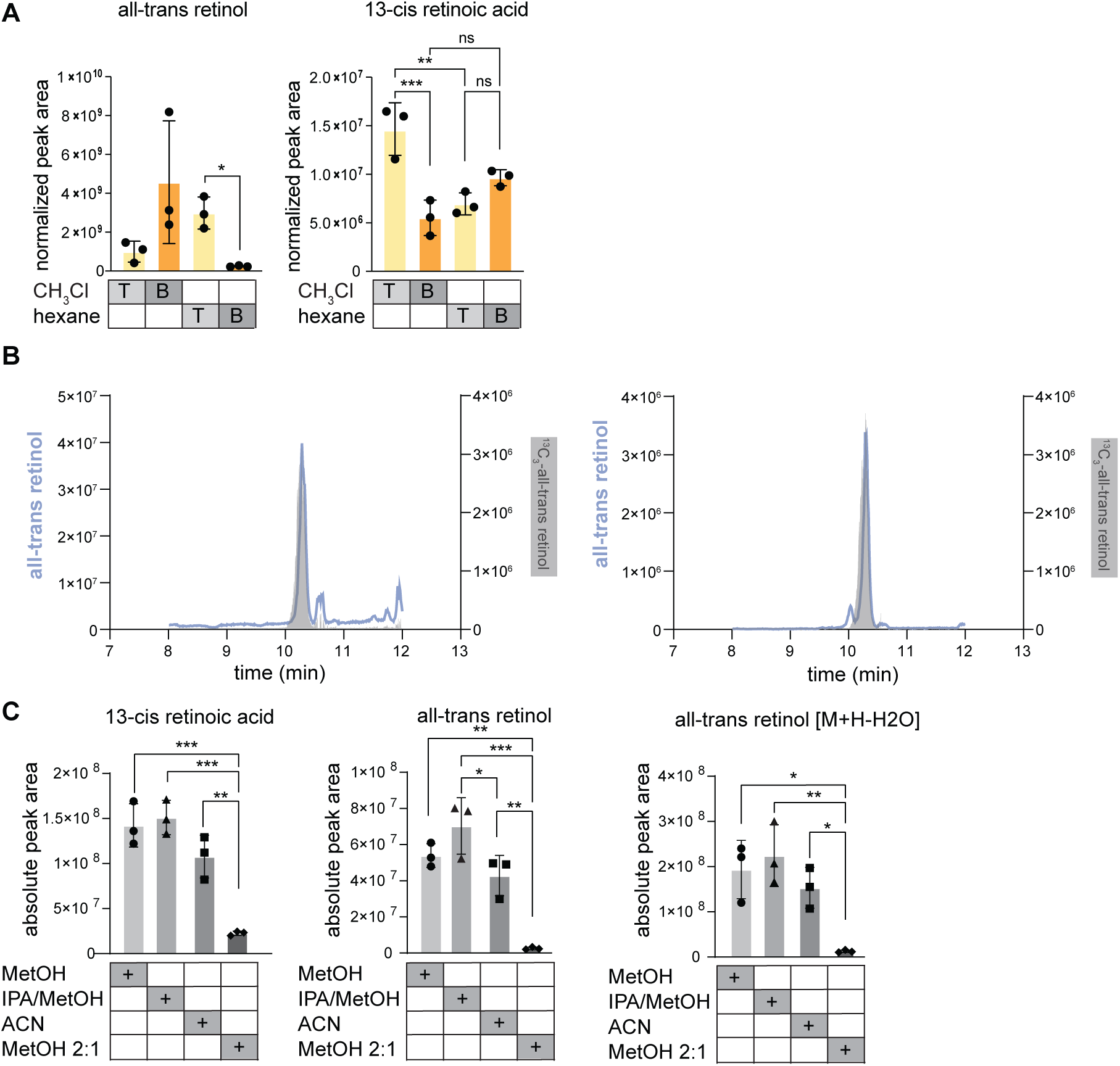
Method optimization of sample preparation for all-trans retinol and 13-cis retinoic acid detection in mouse liver tissue. (A) ^13^C₃-all-trans retinol internal standard normalized peak areas for all-trans retinol and 13-cis retinoic acid in top (labeled as “T”) and bottom (labeled as “B”) phases from two extraction methods in mouse liver tissue. “CH₃Cl” refers to an extraction with a 2:2:1.8 ratio of chloroform:methanol:water, while “hexane” extraction was performed in two steps as detailed in the Methods section. Experiments were repeated 3 times. Statistical significance is calculated by: Welch’s t-test for hexane extraction (left panel) and one-way ANOVA for indicated groups (right panel). (B) Overlay of all-trans retinol and ^13^C₃-labeled internal standard peaks in mouse liver (left) and eye tissue (right) from the top hexane phase. (C) Reconstitution and solubility test. Bar graphs present recovery of indicated retinoids post extraction in 2:2:1.8 chloroform:methanol:water and reconstitution in indicated solvents and mixes. Statistical significance was calculated by one-way ANOVA for indicated groups. Because an internal standard was only added to the extraction buffers, absolute peak areas are shown to enable comparison across extraction methods.

To further assess robustness and generalizability, additional biphasic extraction methods were evaluated across multiple biological and food-derived matrices, including mouse liver, eye tissue, and fish samples (Fig. S5A–D). In mouse liver and eye tissue, chloroform/methanol/water (2:2:1.8), hexanes-based, and isopropanol-based biphasic extractions revealed clear method-dependent differences in retinoid recovery (Fig. S5A). When broad coverage across retinoids was required, chloroform-based extraction provided the most comprehensive recovery, with most compounds detected in the bottom (organic) phase. In contrast, compound-specific preferences were observed for alternative chemistries: all-trans retinol showed highest recovery with hexanes-based extraction, 13-cis retinoic acid and cis/trans retinal were most efficiently recovered using isopropanol-based protocols, and all-trans retinoic acid exhibited the most consistent recovery with chloroform-based extraction. Across all methods, retinoids partitioned between phases to varying extents, indicating that analysis of both phases is advisable for comprehensive profiling. Despite these differences, the 2:2:1.8 chloroform/methanol/water extraction yielded the most consistent bottom-phase recovery across retinoids, supporting its use as a broadly applicable strategy.

Based on this consistency, we next evaluated different chloroform/methanol/water ratios in fish-derived matrices. In cooked and raw salmon and cod samples, chloroform bottom-phase extracts consistently produced higher normalized retinoid signals across all tested solvent ratios (Fig. S5B–D). Among these, ratios of 2:2:1.8 and 2:1:0.5 performed most robustly across matrices, whereas the 4:6:3 ratio showed less consistent performance. Although extraction efficiency varied between experiments and differences did not reach statistical significance (Fig. S5C), similar trends were observed for both 13-cis retinoic acid and retinal. Collectively, these results indicate that while extraction performance depends on matrix and analyte composition, bottom-phase extracts from chloroform/methanol/water-based protocols—particularly the 2:2:1.8 ratio—provide the most consistent recovery across diverse sample types. These results highlight that extraction efficiency and phase preference vary between retinoid species and must be considered when selecting a sample preparation strategy.

To assess reproducibility of the optimized extraction across different tissue types, we compared chromatographic profiles of endogenous and isotope-labeled all-trans retinol extracted from mouse liver and eye tissue (Fig. 3B). All-trans retinol was used as a representative analyte for this comparison. Overlay of endogenous and internal standard signals demonstrated consistent retention times, peak shapes, and relative signal intensities across tissues. These results indicate that the selected extraction protocol yields reproducible retinoid recovery across distinct tissue types, supporting its applicability beyond a single biological matrix. This analysis was intended as a representative validation of reproducibility rather than a comprehensive assessment of matrix effects which could be addressed in extended future work.

Finally, we evaluated the influence of post-extraction reconstitution solvent composition on retinoid signal intensity (Fig. 3C). Extracts were reconstituted in solvents of varying polarity, including pure methanol, pure acetonitrile, methanol/isopropanol mixtures, and methanol-dominant mixtures. More hydrophobic solvent systems (MetOH, IPA/MetOH, ACN) consistently resulted in higher retinoid signal intensities whereas methanol-rich conditions produced reduced recovery (MetOH 2:1 water). Among the tested conditions, the 100% MetOH solvent mixture provided the most robust signal across analytes. These findings demonstrate that reconstitution solvent composition is a critical determinant of retinoid recovery and should be optimized alongside extraction chemistry.

### 3.5 Comparison of extraction efficiency in CSF and liver

To evaluate extraction performance in low-volume, low-abundance matrices, retinoid recovery was compared between mouse cerebrospinal fluid (CSF) and liver using biphasic extraction protocols based on chloroform/methanol/water (2:2:1.8) and hexanes (Fig. S6A–B). The liver serves as the primary storage organ for vitamin A and contains relatively high concentrations of retinoids compared to other tissues(Blaner et al. 2016), Analysis focused on all-trans retinol and 13-cis retinoic acid, including their dominant MS^1^ adducts.

In liver, hexanes-based extraction yielded slightly higher recovery of all-trans retinol, particularly in the first (non-polar) hexanes phase, whereas recovery of 13-cis retinoic acid was modestly higher in the bottom phase. In contrast, retinoids were better detected in CSF samples when the chloroform/methanol/water (2:2:1.8) extraction was used, with higher signals detected in the bottom chloroform phase compared to both the top methanol phase or the hexanes-based extraction fractions, although differences did not reach statistical significance. Finally, these trends were less pronounced for the corresponding isotope-labeled standards, clearly showing that this is largely driven by extraction efficiency rather that solubility per se. While overall extraction trends were broadly similar between liver and CSF, subtle matrix-dependent differences in relative recovery and adduct abundance were observed, supporting the selection of chloroform-based extraction for CSF analyses.

Given the low abundance of retinoids in CSF, we next evaluated whether MS²-based detection could improve analytical sensitivity relative to MS^1^-based quantification. We relied on optimal product ions evaluated in our work with standards (Fig S3 E, F). Signal-to-noise ratios were compared for MS^1^ extracted ion chromatograms and corresponding PRM transitions (Fig. S6C–D). PRM-based detection consistently improved signal-to-noise ratios for retinoids detected at low abundance, with diagnostic fragment ions providing cleaner chromatographic traces and reduced background interference. This improvement was particularly evident in CSF samples, where MS^1^ signals approached the detection limit. These findings demonstrate that PRM-based strategies can enhance confidence in retinoid detection in low-volume, low-concentration biofluids and provide a complementary approach to MS^1^-based quantification.

### 3.6 Retinoid detection in cerebrospinal fluid

Using the optimized chromatographic, mass spectrometric, and sample preparation conditions, we assessed the detectability of retinoids in mouse CSF. All-trans retinol and 13-cis retinoic acid were evaluated using a combination of MS^1^ extracted ion chromatograms and MS²-based PRM quantitation (Fig. 4A–B). All-trans retinol was clearly detected in CSF, with clear chromatographic peaks observed in especially PRM traces. PRM-based detection (diagnostic fragment 93.0702) yielded a signal-to-noise ratio of approximately 5 for the diagnostic fragment ion, providing additional confidence in metabolite identification. Overlay with the isotope-labeled internal standard confirmed consistent retention time and fragmentation behavior. For 13-cis retinoic acid, detection was more variable and approached the lower limits of detection, underscoring the challenges associated with measuring low-abundance retinoids in CSF.

**Figure 4.**
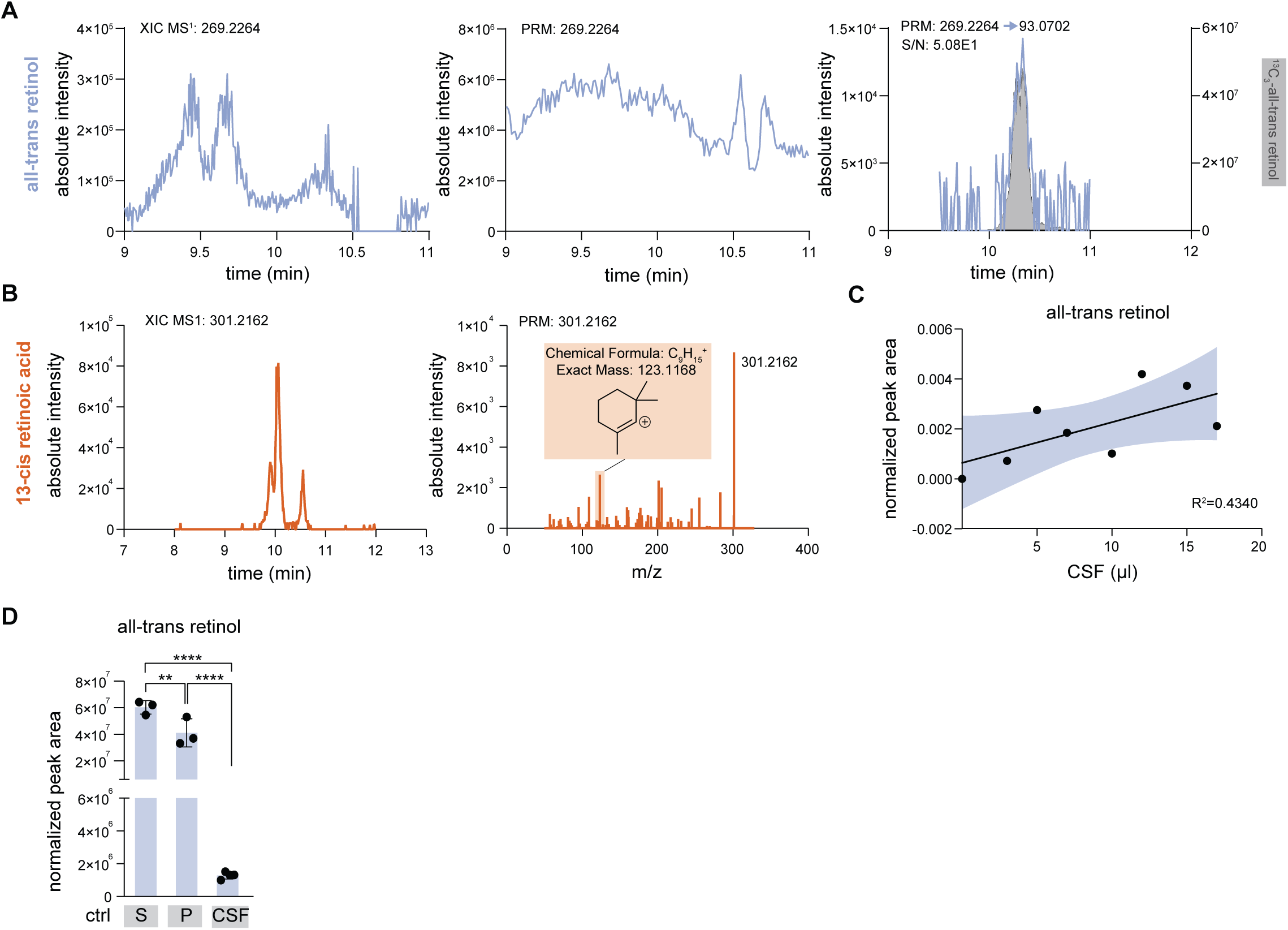
Sample preparation and analysis for retinoid metabolite detection in CSF. (A) XIC (left) of retinol (m/z 269.2264) in mouse CSF and parallel reaction monitoring (PRM) spectrum highlighting the precursor (middle) and diagnostic fragment ion (m/z 93.0702) (right) with internal standard ^13^C_3_-all-trans retinol overlaid. S/N - signal to noise ratio. (B) XIC for 13-cis retinoic acid (m/z 301.2162) in mouse CSF and MS^2^ mass spectra highlighting precursor ion and unique ion fragment (m/z 123.1170). (C) Mouse CSF sample volume optimization for retinoid metabolite detection, extracted with 2:2:1.8 chloroform:methanol:water and resuspended in acetonitrile. Peak areas were normalized to the ^13^C_3_-all-trans retinol internal standard. A linear regression is shown with the corresponding R² and 95% confidence intervals. (D) Bar graphs for serum (“S”), plasma (“P”) or CSF from control mice. One-way ANOVA across all indicated conditions yielded no significant differences. For CSF, an additional Welch’s t-test was performed, with results indicated in parentheses.

To determine minimal sample volume requirements for retinoid detection in mouse CSF, increasing volumes of CSF were extracted and analyzed under identical conditions (Fig. 4C). Despite substantial inter-sample variability, a positive relationship between injected CSF volume and retinoid signal intensity was observed. Linear regression analysis indicated a moderate correlation between CSF volume and all-trans retinol signal (R² = 0.4340), suggesting that increased sample volume improves detectability but does not fully overcome biological or technical variability. These results define practical constraints for CSF-based retinoid analysis in small-volume samples.

Finally, the optimized workflow was applied to compare retinoid levels across serum, plasma, and CSF obtained from control mice (Fig. 4D). All-trans retinol was readily detected in serum and plasma across both groups. In CSF, all-trans retinol was detectable near or at detection limit. These results demonstrate the applicability of the optimized method to measure retinoid levels in low-volume biofluids and the low-volume, low-abundance CSF.

## 4 Conclusions

In this study, we performed an integrated optimization of liquid chromatography, ionization conditions, fragmentation behavior, and extraction procedures for the analysis of retinoids across tissues and biofluids. The results demonstrate that analytical performance is not governed by a single step of the workflow but by the interaction between chromatographic separation, ion formation, and sample preparation. In particular, chromatographic conditions influenced not only retention behavior but also the distribution of detected ionic species, which in turn affected precursor ion selection for MS²-based quantification. Extraction experiments further revealed compound-dependent recovery patterns, while optimized acquisition parameters enabled reliable detection in a low-abundance biofluid, such as the mouse CSF. Together, these findings indicate that accurate retinoid measurement requires coordinated optimization of chromatography, mass spectrometric detection, and extraction, rather than independent adjustment of individual parameters. Earlier targeted LC–MS methods established sensitive retinoic acid quantification in limited samples(Jones et al. 2015; Kane et al. 2005), and our results extend this framework by systematically evaluating how these analytical components influence one another within a single workflow.

The observed analytical behavior reflects intrinsic chemical properties of retinoids. The conjugated polyene backbone allows geometric isomerization, while functional groups such as alcohols, aldehydes, and acids confer partial polarity and chemical reactivity. As a result, retinoids exhibit characteristics of both lipids and small metabolites. These features help explain the column-dependent peak multiplicity, solvent-dependent recovery, and variable ion formation observed in the present study. Retinol readily formed dehydrated ions, whereas retinoic acids displayed multiple adduct species, consistent with their susceptibility to in-source reactions and metal ion association. Such behavior has long complicated retinoid analysis, including sensitivity to light, oxidation, and interconversion during handling(Furr 2004; Ioele et al. 2005). In LC–MS analysis, these chemical properties translate into adsorption losses, variable ionization efficiency, and dependence on solvent composition(Czuba et al. 2020). Our findings suggest that several commonly observed analytical inconsistencies—such as signal variability and multiple peaks for a single analyte—are direct consequences of retinoid chemistry rather than purely instrumental effects.

One of the central observations of this study is that chromatographic conditions influenced not only retention time but also the ionic species detected for individual retinoids. For 13-cis retinoic acid, multiple partially resolved peaks were observed across columns, accompanied by differences in relative adduct abundance. Consequently, precursor ion selection depended on chromatographic configuration. This finding has practical implications because targeted assays frequently monitor a single ion species, yet different chromatographic environments may favor different adducts. Similar considerations have been noted in broader metabolite and lipid analyses, where reliance on a single adduct can affect quantitative accuracy (Czuba et al. 2020). Our results therefore emphasize the importance of MS² confirmation and careful precursor selection when developing targeted retinoid assays. Rather than identifying a universally optimal column, the data demonstrate that chromatographic choice determines which ionic form of a retinoid is quantified and, consequently, how results may compare across studies.

Optimization of ionization and fragmentation parameters showed that retinoids respond in a compound-dependent manner to electrospray source conditions. Variations in S-lens voltage and ion transfer capillary temperature altered signal intensity unevenly across analytes, indicating that uniform acquisition settings may bias detection toward particular retinoids. Fragmentation experiments further demonstrated that intermediate collision energies generated the most reliable diagnostic ions, whereas low energies preserved precursor ions and high energies reduced selectivity through excessive fragmentation. Parallel reaction monitoring was therefore used to confirm analyte identity and quantitation in samples near the detection limit, where MS¹ signals alone were susceptible to background interference. In this context, PRM primarily increased confidence in identification rather than intrinsic sensitivity(Peterson et al. 2012).

Extraction experiments showed that retinoid recovery depended on both analyte identity and sample matrix. Hexanes preferentially recovered retinol species, whereas retinoic acids and retinal species were better recovered with more polar organic phases. Chloroform/methanol/water extraction provided the most consistent overall performance, although individual compounds still partitioned between phases. Importantly, extraction behavior differed between tissues and biofluids, indicating that protocols optimized in tissue cannot be directly transferred to low-protein matrices such as cerebrospinal fluid. These results demonstrate that extraction introduces systematic bias toward particular retinoids and should be selected according to the goals of a study, especially when broad profiling is intended. The sensitivity of retinoid measurements to handling and sample preparation has been recognized previously(Czuba et al. 2020; Furr 2004), and our findings extend this observation by highlighting matrix-dependent recovery across commonly used extraction approaches.

Application of the optimized workflow to cerebrospinal fluid demonstrated that retinoids can be detected in a low-volume, low-abundance biofluid. Signals approached the analytical detection limit, and MS² confirmation was required to distinguish true analyte peaks from background interference. Under these conditions, internal standards and PRM acquisition were particularly important for reliable identification. Previous studies have reported retinol measurements in CSF using targeted approaches(Tabassi et al. 2005), but systematic evaluation of analytical parameters in this matrix has been limited. The present results therefore demonstrate analytical feasibility and provide a framework for future studies investigating retinoids in biofluids rather than establishing biological interpretation.

Several limitations should be considered when applying this workflow. Chromatographic separation of certain retinal isomers remained incomplete under the reversed-phase conditions used here, and adduct variability, particularly for sodium-associated species, may still influence precursor selection in some matrices. In low-abundance samples such as cerebrospinal fluid, signals approached the limit of detection, making MS² confirmation necessary for confident identification. In addition, retinoids are susceptible to light exposure, oxidation, and isomerization during handling, which cannot be fully eliminated despite controlled preparation procedures(Furr 2004; Kane et al. 2005). The present study therefore establishes a workflow suited for relative comparisons across samples rather than a fully validated assay for absolute clinical quantification.

## Author Contributions

Conceptualization: J.B., B.P., N.K. Methodology and investigation: J.B., B.P. Data analysis: J.B., B.P., M.W. Visualization and figure preparation: B.P., J.B. Resources: X.T., A.W., A.B. Writing – original draft: J.B., B.P. Writing – review and editing: N.K. Supervision: B.P., N.K. Funding acquisition: N.K.

## Data availability

Data associated with this study can be requested for the authors and will be made available at MetabolomicsWorkbench: https://www.metabolomicsworkbench.org/

## Acknowledgements

We thank all members of the Kanarek and Lehtinen lab for their advice and help.

