## supplementary information for "Optimization of Retinoid Detection in Cerebrospinal Fluid Using Liquid Chromatography Mass Spectrometry"

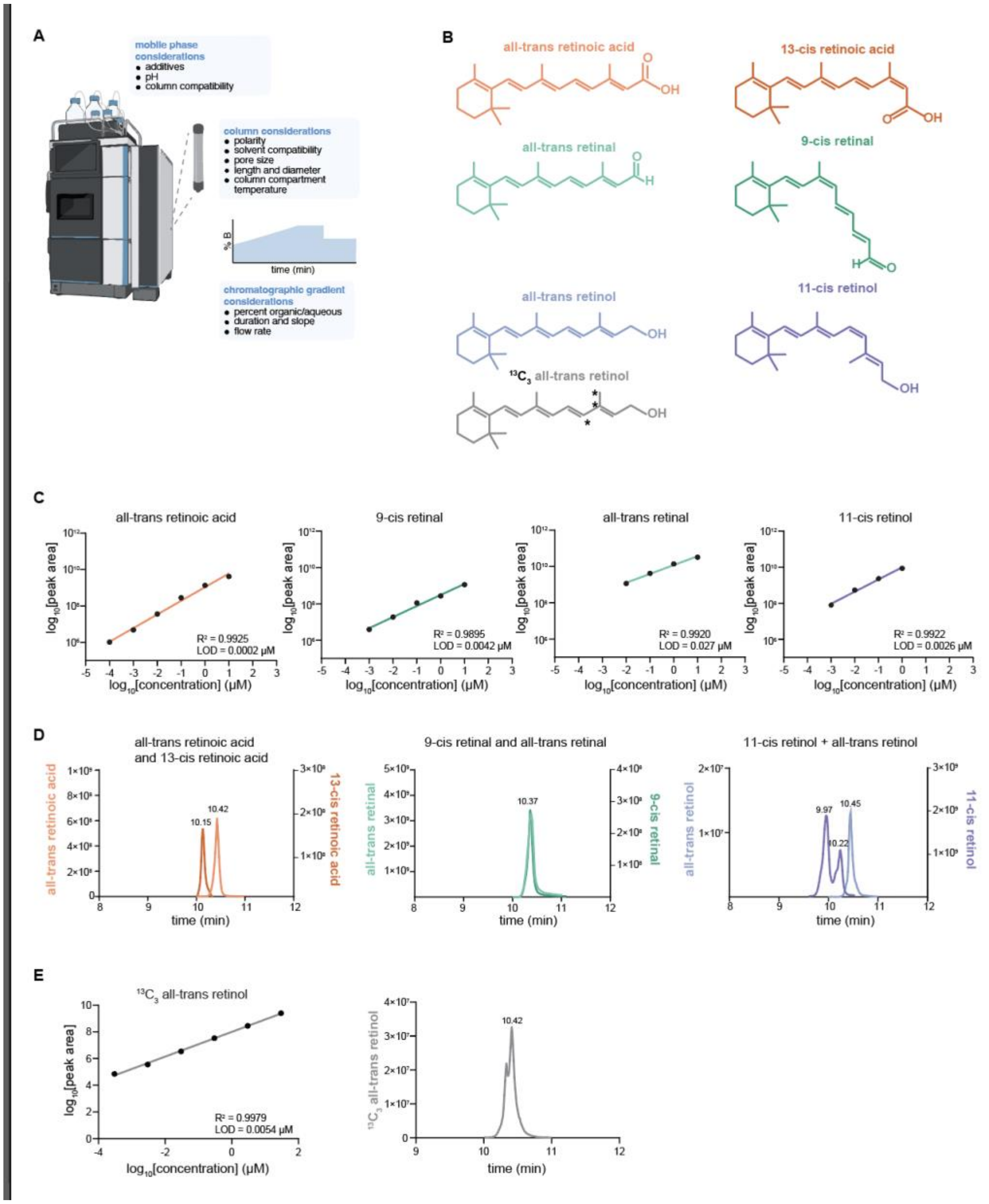


**Figure S1.** Optimization of liquid-chromatography method for the quantification of retinoid metabolites.

**(A)** A scheme describing the method optimization for liquid-chromatography highlighting considerations for mobile phase, column, and chromatographic gradient.

**(B)** Chemical structures of all-trans retinoic acid, 13-cis retinoic acid, all-trans retinal, 9-cis retinal, all-trans retinol, 11-cis retinol, and $^{13}C_3$-all-trans retinol standards.

**(C)**. Limit of detection (LOD) and linearity for indicated retinoids (diluted in methanol) on Ascentis C18-MetOH-2 gradient.

(**D**) Chromatographic peak overlay for indicated retinoids where isomeric compounds are plotted together for comparison of retention times and peak shapes.

(**E**) LOD and linearity for the internal standard $^{13}C_3$-all-trans retinol (diluted in methanol) on Ascentis C18-MetOH-2 gradient.

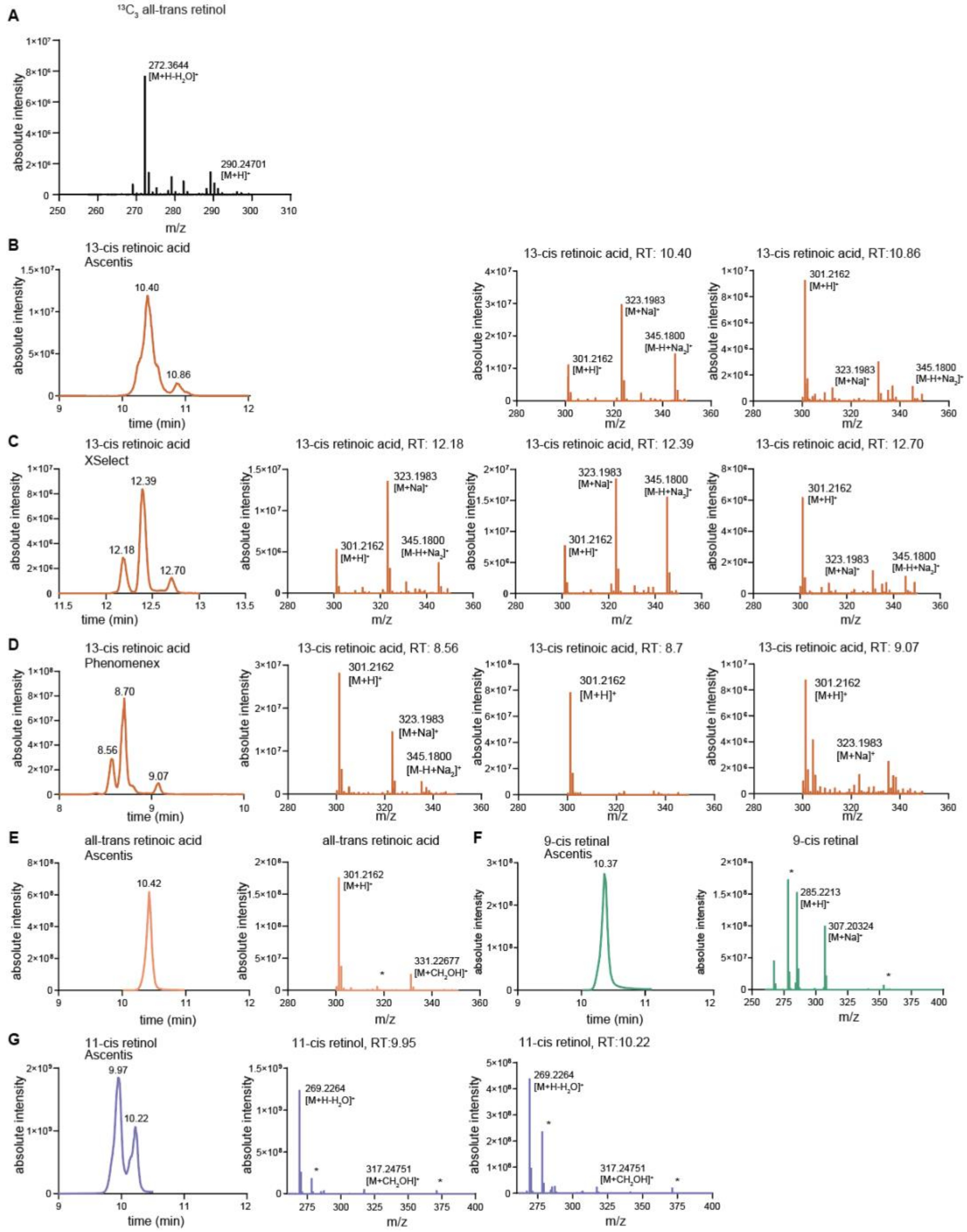


**Figure S2.** Characterization of retention time and adduct abundance for retinoid standards.

(**A**). Abundance of indicated adducts for $^{13}C_3$-all-trans retinol as observed from MS1 spectra.

(**B**, **C, D**) Peak shape and abundance for indicated adducts for 13-cis-retinoic acid on three different C18 columns: Ascentis (**B**), XSelect (**C**), and Phenomenex (**D**) as observed by $MS^1$ spectra. For each isomeric peak, the underlying MS1 spectrum and the most prominent assigned adducts are shown.

(**E, F, G**) As in (**B**) but for all-trans retinoic acid (**E**), 9-cis retinal (**F**) and 11-cis retinol (**G**). Prominent adducts are indicated; asterisks mark co-eluting contaminants.

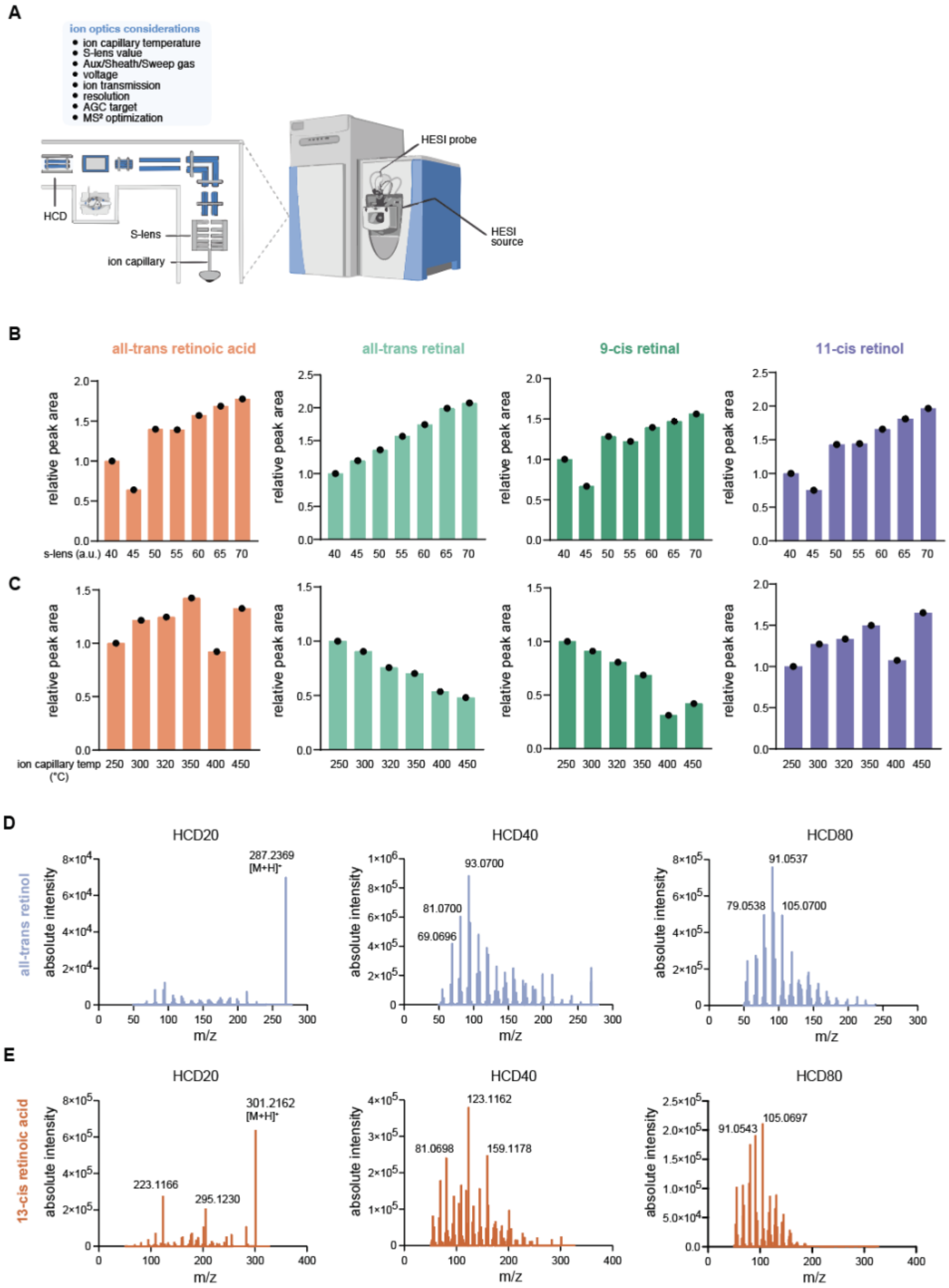


**Figure S3.** Optimization of mass spectrometry method and ionization parameters for the detection of retinoids.

(**A**) Schematic overview of mass spectrometry parameter optimization, highlighting key considerations for HESI ionization settings.

(**B**, **C**) HESI parameter optimization of S-lens (**B**) and ion capillary temperature (**C**) on an Orbitrap mass spectrometer. Bar plots from a representative experiment showing chromatographic peak areas of indicated retinoid metabolites under varying S-lens voltages and ion transfer capillary temperatures.

(**D, E**) Fragmentation pattern under varying collision energies (HCD: 20, 40 or 80 eV) for all-trans retinol (**D**) or 13-cis retinoid acid (**E**) respectively.

A

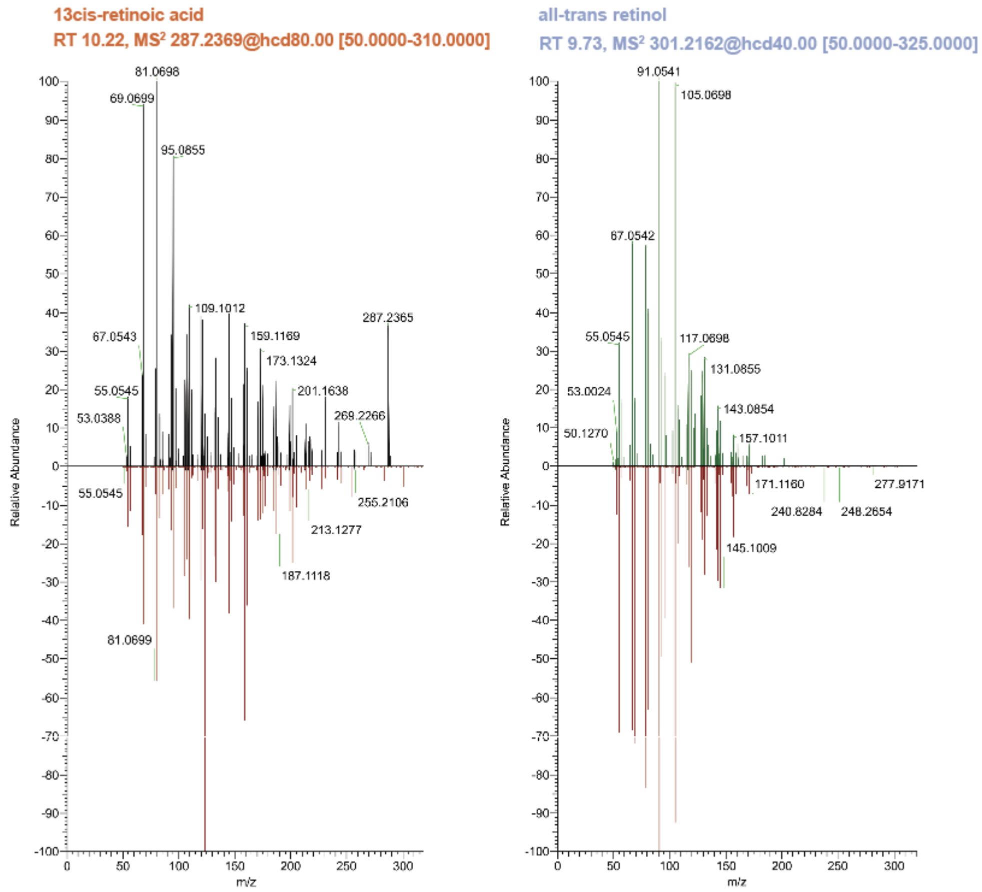


**Figure S4.** Mirror plots between in-house standards (red) and MS$^2$ spectral signal from biological material for the indicated metabolites. Retention times (RT), collision energy (HCD) and individual fragment ion masses are further indicated.

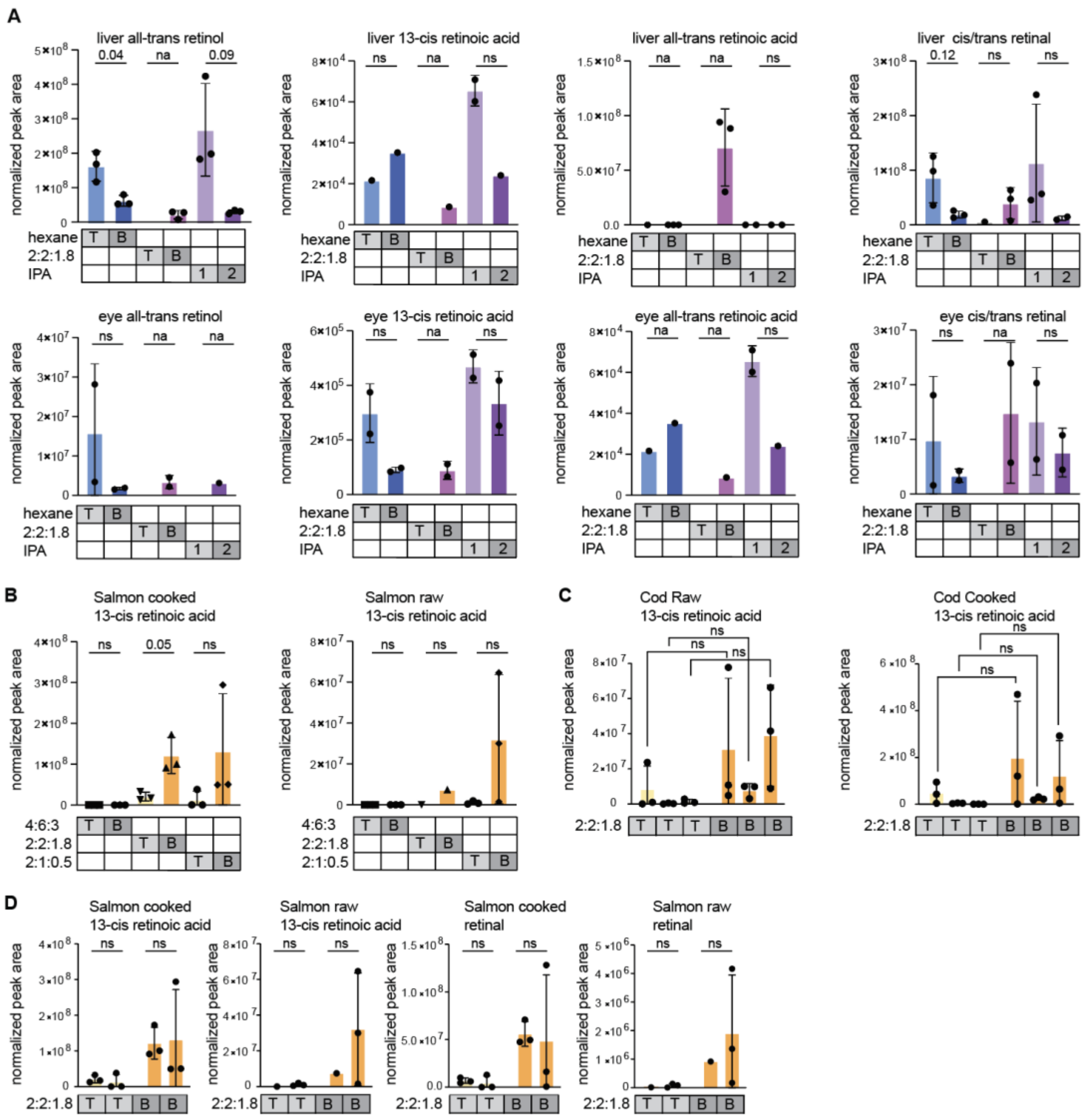

**Figure S5.** Method optimization of the sample preparation for all-trans retinol and 13-cis retinoic acid detection in tissue.

**(A)** Normalized peak areas of retinoids in top (“T”) and bottom (“B”) phases from three extraction methods in mouse liver and eye tissue. The 2:2:1.8 ratio refers to chloroform:methanol:water; hexane and IPA extractions were two-step procedures as described in the Methods section. Peak areas were normalized to internal standards and endogenous metabolites, and across experiments by extraction volume. P-values were obtained by one-way ANOVA with Šídák’s correction for multiple comparisons: * p<0.05, ** p<0.01.

**(B)** Peak areas of retinoid metabolites in top (“T”) and bottom (“B”) phases from three extraction methods in cooked and raw fish. Ratios 4:6:3, 2:2:1.8, and 2:1:0.5 refer to chloroform:methanol:water compositions used in the extractions. p-values were obtained by one-way ANOVA with Šídák’s correction for multiple comparisons: * p<0.05, ** p<0.01.

**(C)** Normalized peak areas for retinoid metabolites in T and B phases of 4:6:3 extraction from three independent experiments between raw and cooked cod liver. Peak areas were normalized within methods to internal standards and endogenous metabolites, and across methods by extraction volume and tissue mass. p-values were obtained by one-way ANOVA with Šídák's correction for multiple comparisons: * p<0.05, ** p<0.01.

(**D**) Normalized peak areas from two independent experiments of 13-cis retinoic acid and retinal in top ("T") and bottom ("B") phases from three extraction methods applied to cooked and raw salmon. Ratios 4:6:3, 2:2:1.8, and 2:1:0.5 indicate the chloroform:methanol:water compositions used in each extraction. Peak areas were normalized to internal standards and endogenous metabolites, and across experiments by extraction volume and tissue mass. p-values were obtained by one-way ANOVA with Šídák's correction for multiple comparisons: * p<0.05, ** p<0.01.

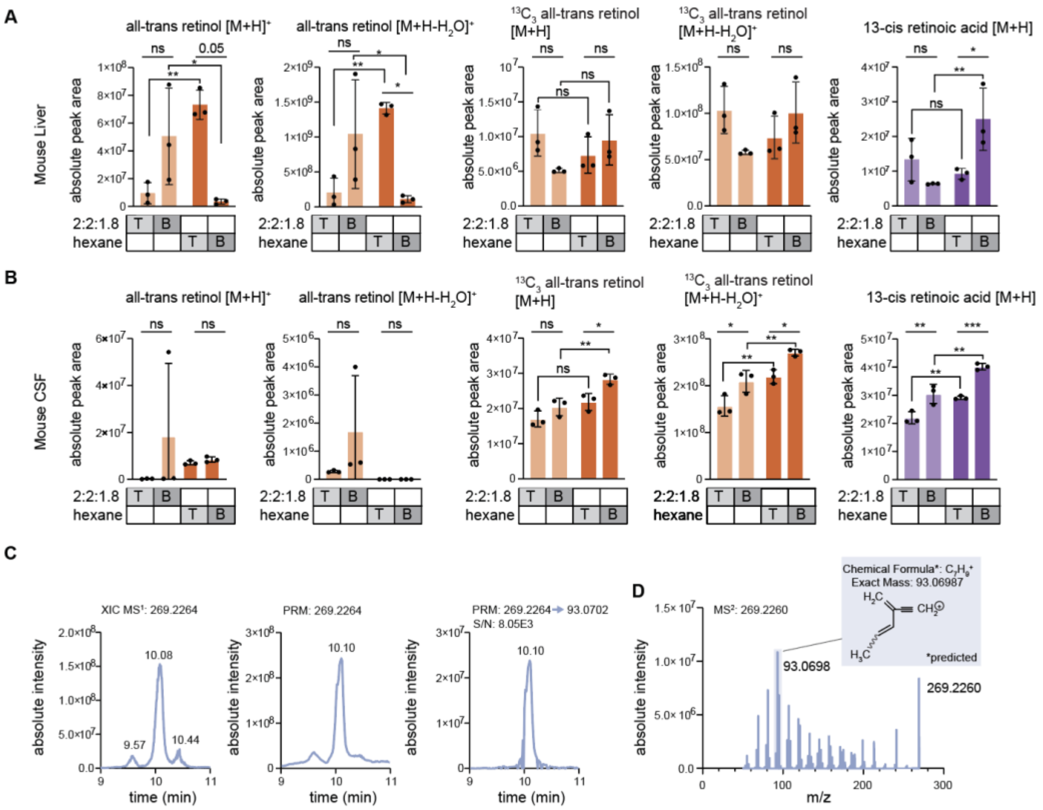


**Figure S6.** Extraction test for retinoids from mouse CSF and liver.

**(A, B)** Normalized peak areas for indicated retinoid adducts and internal standards in top ("T") and bottom ("B") phases from two extraction methods applied for mouse liver (**A**) or CSF (**B**) metabolite profiling. Ratio 2:2:1.8, indicates the chloroform:methanol:water compositions used in the extraction. p-values were obtained by one-way ANOVA with Šídák's correction for multiple comparisons: * p<0.05, ** p<0.01, *** p<0.001.

(**C**) Extracted ion chromatogram (XIC) of retinol (m/z 269.2264) from $MS^1$ (left), parallel reaction monitoring (PRM) chromatogram of the precursor ion (middle), and the diagnostic fragment ion (m/z 93.0702) (right). S/N = signal-to-noise ratio.

(**D**) $MS^2$ spectrum at HCD 40 highlighting the precursor and diagnostic fragment ion (m/z 93.0702) (right). S/N = signal-to-noise ratio.
